# Imaging synaptic density in ageing and Alzheimer’s Disease with [^18^F]-SynVesT-1

**DOI:** 10.1101/2024.10.23.619941

**Authors:** Joseph Giorgio, David N. Soleimani-Meigooni, Mustafa Janabi, Suzanne L. Baker, Xi Chen, Tyler N. Toueg, Robby Weimer, Bastian Zinnhardt, Ari Green, Gil D. Rabinovici, William J. Jagust

**Affiliations:** Department of Neuroscience, University of California Berkeley, Berkeley, California, USA, 94720; School of Psychological Sciences, College of Engineering, Science and the Environment, University of Newcastle, Newcastle, New South Wales, Australia, 2308; Memory and Aging Center, Department of Neurology, Weill Institute for Neurosciences, University of California, San Francisco, San Francisco, CA, USA, 94158; Genentech, Inc, USA; Roche, Switzerland; Lawrence Berkeley National Laboratory, USA; Department of Psychology, Stony Brook University, Stony Brook, New York, USA, 11794

## Abstract

Monitoring synaptic injury in neurodegenerative diseases may provide new insights into the evolution of the degenerative process as well as a potential mechanism to target for preservation of function. Synaptic density imaging with PET is a relatively new approach to this issue. However, there are remaining questions about technical approaches to data analysis including reference region selection, and how specific phenotypic presentations and symptoms of Alzheimer’s Disease (AD) are reflected in alterations in synaptic density.

**Methods:** Using an SV2A PET ligand radiolabeled with the ^18^F isotope ([^18^F]-SynVesT-1) we performed sensitivity analyses to determine the optimal reference tissue modelling approach to derive whole brain ratio images. Using these whole brain images from a sample of young adults, older adults, and patients with varied phenotypic presentations of AD we then contrast regional SV2A density and *in vivo* AD biomarkers.

**Result:** Reference tissue optimisation concluded that a cerebellar grey matter reference region is best for deriving whole brain ratio images. Using these whole brain ratio images, we find a strong inverse association between [^18^F]-SynVesT-1 PET uptake and amyloid beta and tau PET deposition. Finally, we find that individuals with lower temporal grey matter volume but higher temporal [^18^F]-SynVesT-1 PET uptake show preserved performance on the MMSE.

**Conclusions:** [^18^F]-SynVesT-1 PET shows a close association with *in vivo* AD pathology and preserved SV2A density may be a possible marker for resilience to neurodegeneration.

## Introduction

Alzheimer’s Disease (AD) is characterised by the presence of amyloid beta (Aβ) and tau deposits each of which has a profound effect on the number of functional synapses in the brain (*1*). *In vitro* approaches to quantify the underlying density of synapses have involved measurements of pre-synaptic proteins such as synaptophysin and synaptic vesicle glycoprotein 2A (SV2A), with a close association observed between SV2A and synaptophysin (*2*). Recently, PET ligands derived from the structure of levetiracetam, an antiepileptic drug that selectively binds to SV2A, were created to measure *in vivo* SV2A. The most widely utilised SV2A PET ligand is [^11^C]-UCB-J, which shows high brain uptake, fast kinetics, high test-retest performance, and can be displaced by administering levetiracetam (*2–4*). However, clinical application of [^11^C]-UCB-J is inherently limited due to the short half-life of the ^11^C isotope. Subsequent SV2A PET ligands radiolabeled with the ^18^F isotope have been developed, with [^18^F]-SynVesT-1 (also known as [^18^F]-SDM8) shown to have favourable PET qualities, with excellent kinetic and *in vivo* binding properties, and high test-retest reproducibility (*5*,*6*).

*In vivo* SV2A PET in healthy young adults has suggested the highest cortical synaptic density in the posterior cingulate, precuneus, and temporal cortex with these relative differences also observed through *in vitro* autoradiography (*7*). Within subcortical regions, the highest density was observed in the striatum, with lower density in the hippocampus and cerebellum (*7*). These patterns of high synaptic density do not appear to be correlated with regions of high cortical thickness or subcortical volume (*7*,*8*). SV2A PET has been deployed to study synaptic variations due to ageing (*8–10*), non-AD related neurodegenerative disorders (*11–22*), and psychiatric disorders (*23–25*). In addition, SV2A PET has been shown to be lower in patients with AD compared to controls and is associated with variations in cognition, glucose hypometabolism, Aβ PET, and tau PET (*26–36*), however only one of these research groups utilised [^18^F]-SynVesT-1 (*36*). These prior studies into AD typically investigated amnestic presentations of AD or MCI and it remains to be seen how SV2A PET binding varies across non-amnestic or atypical AD phenotypes.

To relieve the burden on participants, non-invasive reference tissue modelling of PET data is preferred. Initial studies on optimal reference regions for [^11^C]-UCB-J PET selected the centrum semiovale within the cortical white matter due to the low concentration of SV2A compared to grey matter regions (*2*,*37*). Although the centrum semiovale is the most frequently used reference region in SV2A PET studies, using the cerebellum as a reference region is becoming more common (*38*,*39*). This is pressing as recent studies utilising reference tissue modelling of [^18^F]-SynVesT-1 in neurodegenerative disorders have used different reference tissues (white matter: (*40*); cerebellum: (*36*)), this is despite ground truth arterial sampling methods indicating the white matter is optimal as a reference tissue (*6*). Therefore, direct comparison of simplified reference tissue modelling using cerebellar and white matter reference regions across clinically impaired, young, and older populations is required to determine the optimal reference tissue for modelling [^18^F]-SynVesT-1 PET uptake.

In this study we deployed [^18^F]-SynVesT-1 PET in a sample of young and cognitively normal older adults as well as clinically impaired patients with varied AD phenotypic presentations. Using full dynamic [^18^F]-SynVesT-1 acquisitions, we tested how well simplified reference tissue model fits captured cortical time activity curves for various reference regions. After determining the optimal reference region, we derived summary images to investigate variation in [^18^F]-SynVesT-1 PET uptake across ageing and AD phenotypes. We then tested associations between [^18^F]-SynVesT-1 PET uptake, Aβ PET, tau PET, and cognitive impairment. Finally, we evaluated whether there is additional information in [^18^F]-SynVesT-1 PET compared to MRI volumetric measures. We hypothesise that SV2A uptake will be reduced in patients compared to controls and that patients with atypical AD presentations will have patterns of decreased SV2A uptake representative of the AD phenotype.

## Methods

### Participants

In this study, we acquired dynamic [^18^F]-SynVesT-1 PET imaging on 28 participants (2 participants had incomplete scanning sessions). Participants were selected from three populations recruited from two independent sites. 7 young adults (Y) and 14 cognitively normal older adults (O) were recruited through the Berkeley Ageing Cohort Study (BACS). 7 patients (P) with clinical presentations of Alzheimer’s disease (AD) were recruited from University of California San Francisco (UCSF) Memory and Ageing Center. These patients had a mix of phenotypic presentations: amnestic dementia (n=3), logopenic variant of primary progressive aphasia (lvPPA) (n=1), posterior cortical atrophy (PCA) (n=1), amnestic mild cognitive impairment (MCI) (n=1), and non-amnestic MCI (n=1). 43% of older adults and 86% of patients had visually positive/elevated Aβ-PET scans (1 amnestic MCI was Ab-) (**Table 1**). Y participants did not undergo Aβ or tau PET. This study was approved by the relevant Institutional Review Boards of the University of California, Berkeley and San Francisco and the Lawrence Berkeley National Laboratory (LBNL).

**Table 1.** Participant Demographics.

|  | **Young Adults** | **Older Adults** | **Patients** |
| --- | --- | --- | --- |
| **Sample size** | 7 | 14 | 7 |
| **Age (mean±std)** | 26.1±3.8 | 80.2±4.4 | 66.4±7.5 |
| **Sex (F)** | 4 | 8 | 2 |
| **Race (White)** | 3 | 13 | 4 |
| **Education (mean±std)** | 19±1.6 | 17.4±1.3 | 17.5±2.8 |
| **MMSE (mean±std)** | 29±1.2 | 28.6±1.5 | 23.6±9 |
| **Aβ+** | NA | 6 | 6 |
| **AD tau+** | NA | 4 | 5 |

### Structural MRI

#### BACS participants

Structural MRIs were collected on either a 1.5T MRI scanner at LBNL or a 3T MRI scanner at UC Berkeley including 3D T1-weighted whole brain magnetization prepared rapid gradient echo (MPRAGE) sequences. The 1.5T MPRAGE acquisition parameters were: repetition time (TR) = 2110 ms, echo time (TE) = 3.58 ms, flip angle = 15°, voxel size 1 mm isotropic. The 3T MPRAGE acquisition parameters were: TR = 2300ms, TE = 2.98ms, flip angle = 9°, voxel size 1mm isotropic.

#### UCSF patients

Structural MRIs for UCSF participants were acquired on a 3T Siemens Prisma Fit scanner at UCSF using T1-weighted MPRAGE sequence: TR= 2300 ms, TE = 2.9 ms, flip angle = 9°, voxel size 1 mm isotropic.

All Structural T1-weighted images were analysed using the same processing pipeline. Images were first segmented into grey matter, white matter, and CSF components in native space using Statistical Parametric Mapping (SPM12). Native space T1 images were additionally parcellated with Freesurfer v.5.3.0 using the Desikan-Killany atlas (*41*). These regions of interest (ROIs) were used to define uptake for all PET tracers.

### Ab PET

#### BACS participants

Ab PET imaging of BACS older participants involved injection of ∼15[mCi of [^11^C]-PiB tracer into an antecubital vein, and dynamic acquisition frames were obtained over a 90 min interval on a Siemens Biograph PET/CT (4 × 15 s frames, 8 × 30 s frames, 9 × 60 s frames, 2 × 180 s frames, 8 × 300 s frames, and 3 × 600 s frames) following a head CT. Distribution volume ratios (DVRs) were generated with Logan graphical analysis (*42*) on the aligned PiB frames using the native-space cerebellar grey matter as a reference region calculating the slope from 35-90min post-injection.

#### UCSF patients

All UCSF participants had [^18^F]-Florbetaben (FBB) Aβ PET scans acquired on a Siemens Biograph 64 Vision 600 PET/CT. FBB was administered via intravenous injection of ∼8 mCi of FBB, followed by acquisition of 4 × 5 minute PET frames at 90-110 minutes post-injection. PET frames were realigned, averaged, and co-registered onto their corresponding MRI. Standardized uptake value ratio (SUVR) images were created in native space using an MRI-defined whole cerebellar reference region.

For each participant, a global Aβ index was derived using the centiloid (CL) scale (*43*). CL values were derived for each cohort independently using the GAAIN reference data and cohort specific pipelines and conversion equations (https://www.gaain.org/centiloid-project) (*44*,*45*). Scans were also visually classified as elevated (positive) or non-elevated (negative) using validated, tracer-specific criteria by a neurologist expert reader (DNSM) (*46*).

### Tau PET

#### BACS participants

The [^18^F]-Flortaucipir (FTP) PET protocol entailed the injection of ∼10[mCi of tracer followed by acquisition of PET images 80–100 min post injection on a Siemens Biograph PET/CT scanner.

#### UCSF patients

UCSF participants underwent either FTP (n=3) or [^18^F]-PI-2620 PET (n=4) on the Siemens Biograph 64 Vision 600 PET/CT. The FTP PET protocol involved injection of ∼10 mCi of FTP followed by the acquisition of PET images 75-105 min post injection. The PI-2620 protocol involved the injection of ∼5 mCi of [^18^F]-PI-2620 followed by the acquisition of PET images 30-75 min post injection, of which the 45-75 min data was used here.

For all tau PET scans, PET frames were realigned, averaged, and co-registered onto their corresponding MRI. SUVR images were created in native space using an MRI-defined inferior cerebellar grey matter reference region.

### [^18^F]-SynVesT-1 PET acquisition

[^18^F]-SynVesT-1 was synthesized at the Biomedical Isotope Facility (BIF) at LBNL using a modified iPhase MultiSyn synthesis module. The ^18^F was produced using a Siemens Eclipse (RDS111) medical cyclotron. The tin precursor (Me_3_Sn-SDM-8) in N,N-dimethylacetamide (DMA) was used to carry out the incorporation reaction of ^18^F. After several steps (purification, isolation and reformulation), the desired tracer ([^18^F]-SynVesT-1) was afforded in a solution of ethanol/saline (5% / 95%, respectively). Typical molar activity ranged between 3000-7000 mCi/umol.

5 mCi of [^18^F]-SynVesT-1 was injected, and data were collected on the Siemens Biograph PET/CT scanner for 90 min across 35 dynamic frames (4 x 15, 8 x 30, 9 x 60, 2 x 180, 10 x 300, and 2 x 600s), two O did not complete the full acquisition (33 and 34 frames). A CT scan was performed preceding the PET acquisition for attenuation correction. All PET images were reconstructed using an ordered subset expectation maximization algorithm, with attenuation correction, scatter correction, and smoothing with a 4mm Gaussian kernel. An average of frames within the first 20 min was used to calculate the transformation matrix to co-register the [^18^F]-SynVesT-1 images to the participants’ structural MRI. This transformation matrix was then applied to all aligned [^18^F]-SynVesT-1 frames.

### Optimising modelling of [^18^F]-SynVesT-1 PET

#### Reference region selection

To assess the optimal reference region to use when deriving whole-brain SUVR images, we searched across 3 *a priori* reference regions taken from the native space including the cerebellar grey matter, whole cerebellum, and eroded white matter. The cerebellar reference regions were taken from the Desikan-Kiliany ROIs in native space, and the eroded white matter reference region was defined by smoothing the white matter segmentation from the Desikan-Kiliany ROIs in native space, applying an 8mm^3^ smoothing kernel, and selecting voxels within the Desikan-Kiliany ROIs with a value greater than 0.8. This eroded white matter region encompasses the centrum semiovale (*47*) and was favoured as a white matter reference region due to the relative ease of mask derivation and relative homogeneity in white matter signal.

First, to assess if there was systematic variability in tracer binding within these regions, we extracted standardised uptake values (SUVs) in each reference region from the averaged 70-90-minute frames. In addition to assessing the variability within each reference region based on the coefficient of variation (CV: ratio of population standard deviation and mean), we contrasted SUV values based on diagnostic category and age. Second, we used the simplified reference tissue model 2 (SRTM2) (*48*) to determine how fits of estimated time activity curves (TAC) and tracer clearance from the reference tissue (k_2_’) varied for each reference ROI. We ran SRTM2 using in house MATLAB functions, performing a grid search to determine the optimal parameters to minimise the sum squared error (SSE) between model derived and observed TAC. For computational efficiency and stability, we performed the fitting on the 34 bilateral cortical Desikan-Killiany ROIs rather than on the voxel level. We then averaged the SSE across cortical ROIs to allow us to compare average SSE and k_2_’ across reference regions. From the derived values above, we identified an optimal reference region defined by a low average SSE between observed and fit TAC as well as a low CV of SSE, k_2_’, and SUV. In addition, we required no, or negligible, relationships between age or diagnostic category and model extracted values (SSE between observed and fit TAC, SSE, k_2_’, and reference region SUV).

#### Deriving ratio images

After determining the optimal reference region, we used the group-averaged k_2_’ values to derive whole brain distribution volume ratio (DVR) images with Logan graphical analysis over 35-90 min of data (*42*). We confirmed the decision to use this window by assessing if the tracer was in steady state. To determine this, we used Logan graphical analysis on the average TAC from all cortical grey matter voxels, calculating the slope of Logan X and Logan Y values over a 35-90 min, and 70-90 min post injection window of data. We then compared the similarity of these slopes to understand if there were non-linearities in the emission data that may affect the derivation of DVR values through Logan graphical analysis.

The whole brain DVR images were then used as the benchmark values when assessing the optimal post injection acquisition windows to derive SUVR images. To determine the optimal scanning window, we derived whole brain SUVR images across different 20-minute time windows for participants with 90-minutes of full dynamic acquisitions. We then extracted average uptake within the bilateral Desikan-Killiany cortical ROIs from the DVR image and three SUVR images derived using different 20-minute windows (50-70 min, 60-80min, 70-90min). To determine the validity of each window, we compared ROI uptake in each of the three SUVR images and the DVR image. First, we calculated the shared variance between the ROI values, treating each ROI for each subject as an observation. Second, we assessed the within subject differences in shared variance, treating the within subject shared variance between ROI uptake as a within subject repeated observation.

#### Partial Volume Correction

We used a 2-tissue compartment, partial volume correction (PVC) approach (*49*). To do this, we used in house MATLAB and SPM12 functions to combine grey and white matter binary masks in native space, which were subsequently smoothed using a kernel based on the resolution of the LBNL Biograph PET scanner that collected the [^18^F]-SynVesT-1 PET data (full width at half maximum=[6.5 6.5 7.25]). Each voxel within each frame of the PET data was divided by the value within the same voxel in this smoothed tissue compartment to increase signal on the border of grey matter and CSF. This PVC approach will account for potential partial volume effects predominantly driven by atrophy differences. We then recalculated the optimal k_2_’ using SRTM2 on each PVC frame and derived a PVC DVR image based on the group average k_2_’.

### Associating [^18^F]-SynVesT-1 uptake with clinical and imaging variables

We extracted average tracer uptake from the DVR and SUVR images in four summary regions of interest (ROI), including the meta-temporal ROI (volume weighted average of the entorhinal, amygdala, parahippocampal, fusiform, inferior temporal, and middle temporal ROIs) (*50*), temporal lobe, parietal lobe, and frontal lobe. We *a-priori* selected the meta-temporal ROI as it has shown to be sensitive to AD related changes in volume (*51*) and pathology (tau) (*50*), our limited sample size precluded more exploratory regional analyses. We then contrasted the average uptake within these regions for each of the diagnostic groups (Y, O, P). Next, we assessed the topographic association between [^18^F]-SynVesT-1 and tau PET uptake using the signal from 34 left and 34 right cortical (68 total) Desikan-Killiany ROIs in each participant’s set of [^18^F]-SynVesT-1 and tau PET images. For each participant, we calculated the Pearson’s correlation coefficient between the tau and [^18^F]-SynVesT-1 PET uptake in these 68 ROIs. We then used this correlation coefficient as a measure of the topographic relationship between tau and [^18^F]-SynVesT-1 PET uptake for each participant. We used the PVC [^18^F]-SynVesT-1 DVR image in this analysis, as the two-tissue compartment correction will account for within-subject topographic variation in atrophy that could confound the spatial association between [^18^F]-SynVesT-1 and tau PET uptake. As participants were scanned using multiple tau PET ligands, this precluded us from analysing associations between [^18^F]-SynVesT-1 and tau PET uptake at the group level.

Next, we tested the association between Aβ levels and global cortical [^18^F]-SynVesT-1 uptake. To do this, we derived Aβ centiloid (CL) values using the standard CL region and approach defined in Klunk et al. (*43*). We then extracted the volume weighted average of [^18^F]-SynVesT-1 uptake within Desikan-Killiany cortical ROIs to derive a cortical average of [^18^F]-SynVesT-1 uptake and assessed the association between Aβ load and cortical [^18^F]-SynVesT-1 uptake.

Finally, we tested if individual variability in [^18^F]-SynVesT-1 uptake was associated with cognitive performance on the mini-mental state examination (MMSE). In particular, we assessed if there was additive information in [^18^F]-SynVesT-1 uptake when considering the effect of grey matter volumetric differences in predicting cognition. We evaluated this using multiple linear regression to model the effects of the meta-temporal ROI [^18^F]-SynVesT-1 uptake, meta-temporal ROI grey matter volume (normalised by total intracranial volume), and their interaction on MMSE, including age as a confounding variable. Unless otherwise stated all analyses were repeated using both PVC and non-PVC data.

## Results

### Selecting the optimal reference region and acquisition window for reference tissue images

For the 26 participants who completed 90-minute full dynamic [^18^F]-SynVesT-1 PET acquisition, we did not observe a significant relationship between SUV and age in any reference region (Pearson’s correlation: cerebellar grey matter r(24)=0.22 p=0.3; whole cerebellum r(24)=0.23 p=0.28; eroded white matter r(24)=0.24 p=0.26, one outlier was removed), nor were there statistical differences in SUVs in any reference region when comparing the Y, P, and O groups (one-way ANOVA: cerebellar grey F(2,23)=0.95 p=0.4; whole cerebellum F(2,23)=0.99 p=0.39; eroded white matter F(2,23)=1.47 p=0.25) (**Figure 1**). All reference regions had similar CV (0.237-0.238), although the uptake in the eroded white matter was lower than the two cerebellar reference regions for all but one participant (cerebellar grey matter [mean±std], 4.4±1.04; whole cerebellum, 4.1± 0.97; eroded white matter, 1.62± 0.38).

**Figure 1.**
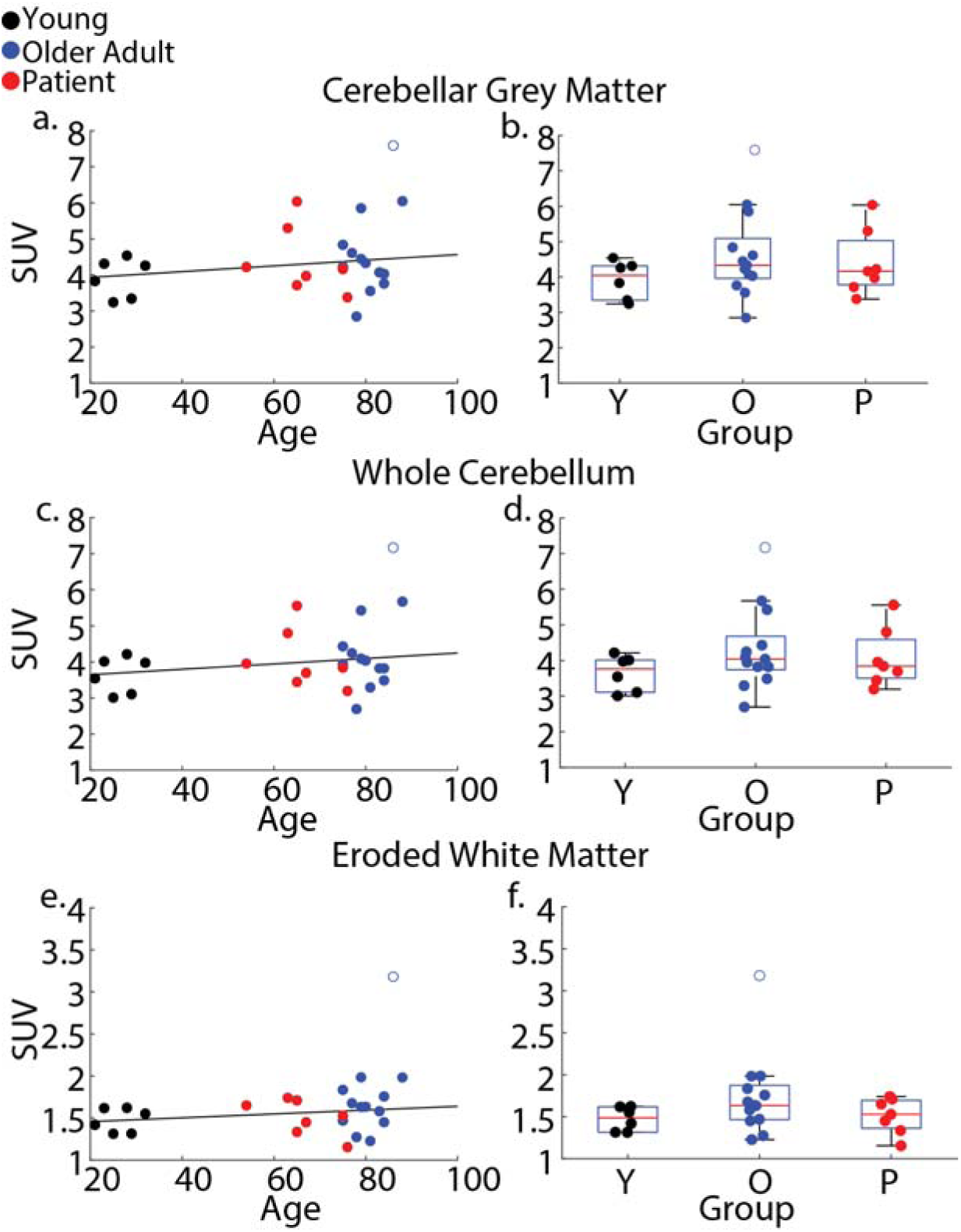
Standardised Uptake Values (SUV) in different reference regions. The left panel shows the association between age and SUV for the (a.) cerebellar grey matter, (c.) whole cerebellum, and (e.) eroded white matter, the black line shows the least squares best fit between age and SUV. The right panel shows distributions of SUV in each of the participant groups young controls (Y), old controls (O), and patients (P) from the (b.) cerebellar grey matter, (d.) whole cerebellum, and (f.) eroded white matter. The unfilled blue data point is an outlier O who is excluded when fitting age and SUV in the different reference regions. Black, blue, and red data points are for young (Y), older adults (O) and patients (P), respectively.

All k_2_’ values in all 28 participants were reliably estimated for each reference region with low CV amongst the sample (**Supplementary Table 1**). Furthermore, cortical TAC were fit with similar accuracy to observed TAC among each group with no significant differences in SSE between Y, P, or O groups (one-way ANOVA: cerebellar grey matter F(2,25)=0.7 p=0.51; whole cerebellum F(2,25)=0.52 p=0.6; eroded white matter F(2,25)=1.7 p=0.20). Similarly, we did not observe an effect of age on TAC fit (Pearson’s correlation: cerebellar grey matter r(24)=0.22 p=0.3; whole cerebellum r(24)=0.23 p=0.28; eroded white matter r(24)=0.24 p=0.26) (**Figure 2**). However, using the cerebellar grey matter reference region, the fit of the SRTM2-estimated TAC with the observed TAC showed the least error (one-way repeated measures ANOVA: F(2,54)=32.4, p<0.0001; cerebellar grey matter vs whole cerebellum ΔSSE=-66304, p<0.0001, cerebellar grey matter vs eroded white matter ΔSSE=-139630, p<0.0001, Tukey HSD corrected) (**Figure 2**). Taken together, these results indicate that a cerebellar grey matter reference region is preferred when deriving DVR images. Cerebellar grey matter will be used as the reference region throughout subsequent analyses. Next, for the 26 participants with full emission data we extracted slopes from Logan graphical analysis using different sampling windows. We observed highly similar slopes when fitting the data in the 35-90 minute or the 70-90 minute windows (**Supplementary Figure 1**.), although the slope using the 70-90 minute window was slightly lower (slope 35-90 min -70-90 min = 0.01; t(25)=2.49, p=0.02). As a difference of 0.01 is only 2.5% of the dynamic range within our sample we selected the 35-90 min window for the derivation of DVR values.

**Figure 2.**
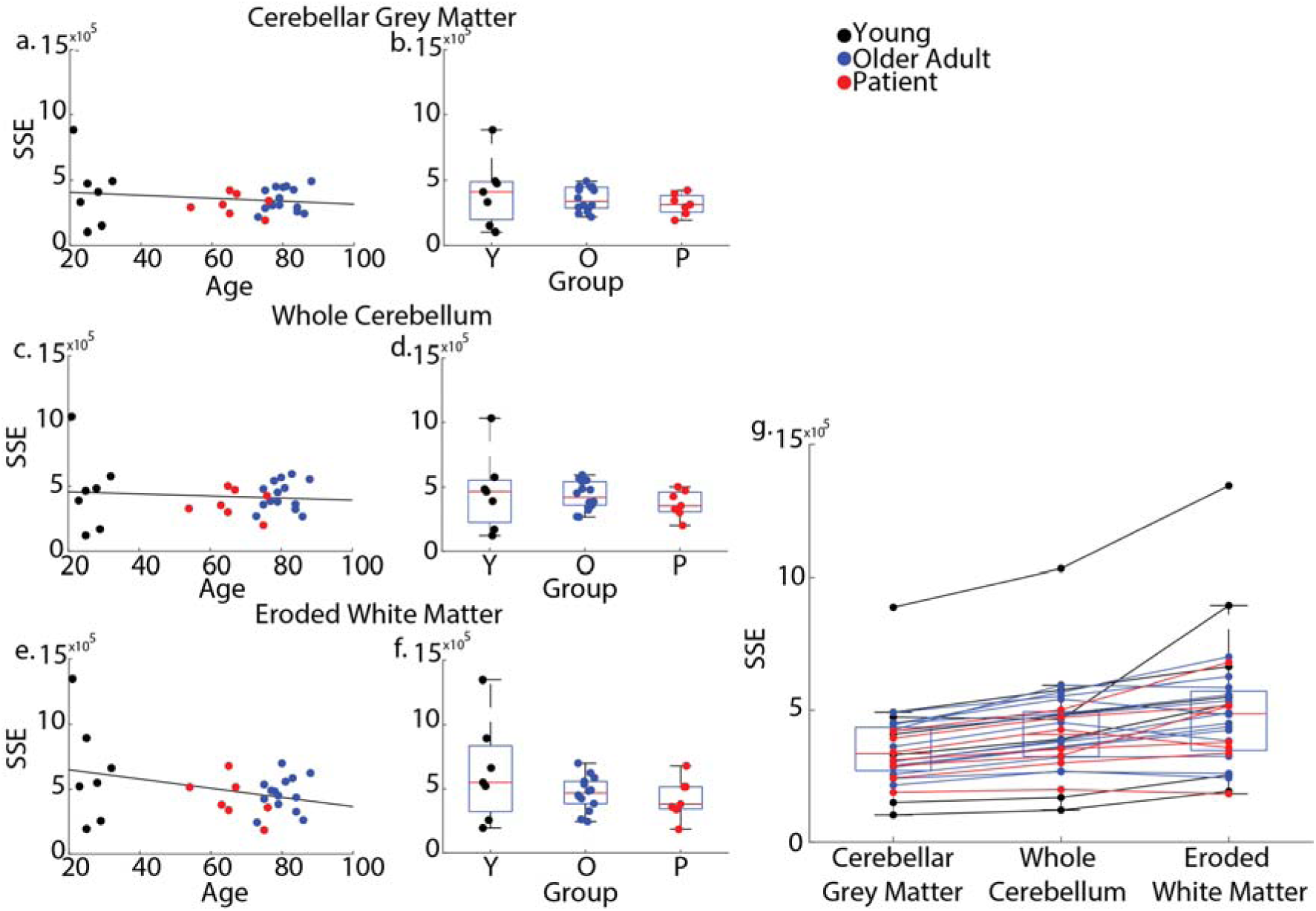
SRTM2 goodness of fit for different reference regions. Relationship of sum squared error (SSE) of SRTM2 predicted time activity curves and observed time activity curves with age using a (a.) cerebellar grey matter, (c.) whole cerebellum, or (e.) eroded white matter reference region. Relationship of SSE of SRTM2 predicted time activity curves and observed time activity curves with participant group using a (b.) cerebellar grey matter, (d.) whole cerebellum, or (f.) eroded white matter reference region. (g.) The within subject SSE of SRTM2 predicted time activity curves and observed time activity curves using a cerebellar grey matter, whole cerebellum, or eroded white matter reference region. Black, blue and red data points are for young (Y), older adults (O), and patients (P), respectively.

We observed high shared variance (R^2^) across all subjects and ROIs between DVR and the different SUVR windows, with the highest occurring in the 60-80 min acquisition window (50-70 min R^2^=0.949; 60-80 min R^2^=0.965; 70-90 min R^2^=0.959). Furthermore when calculating the slope of the least squares fit between DVR and SUVR we observed a slope closest to 1 (slope=0.991) in the 60-80 min window (**Figure 3**). Comparing the percentage of shared variance within subject showed significant differences between the windows, with the closest association between DVR and SUVR values observed using a 60-80 min acquisition window (one-way repeated measures ANOVA: F(2,50)=3.84, p=0.028; 60-80 vs 50-70 ΔR^2^=1.6%, p=0.006, 60-80 vs 70-90 ΔR^2^=1.9%, p=0.011, Tukey HSD corrected) (**Figure 3**). Taken together, these results indicate that an SUVR image derived using emission data 60-80 min post-injection is highly representative of data extracted from the full dynamic acquisition. 60-80 min SUVR is used in all the subsequent SUVR analyses.

**Figure 3.**
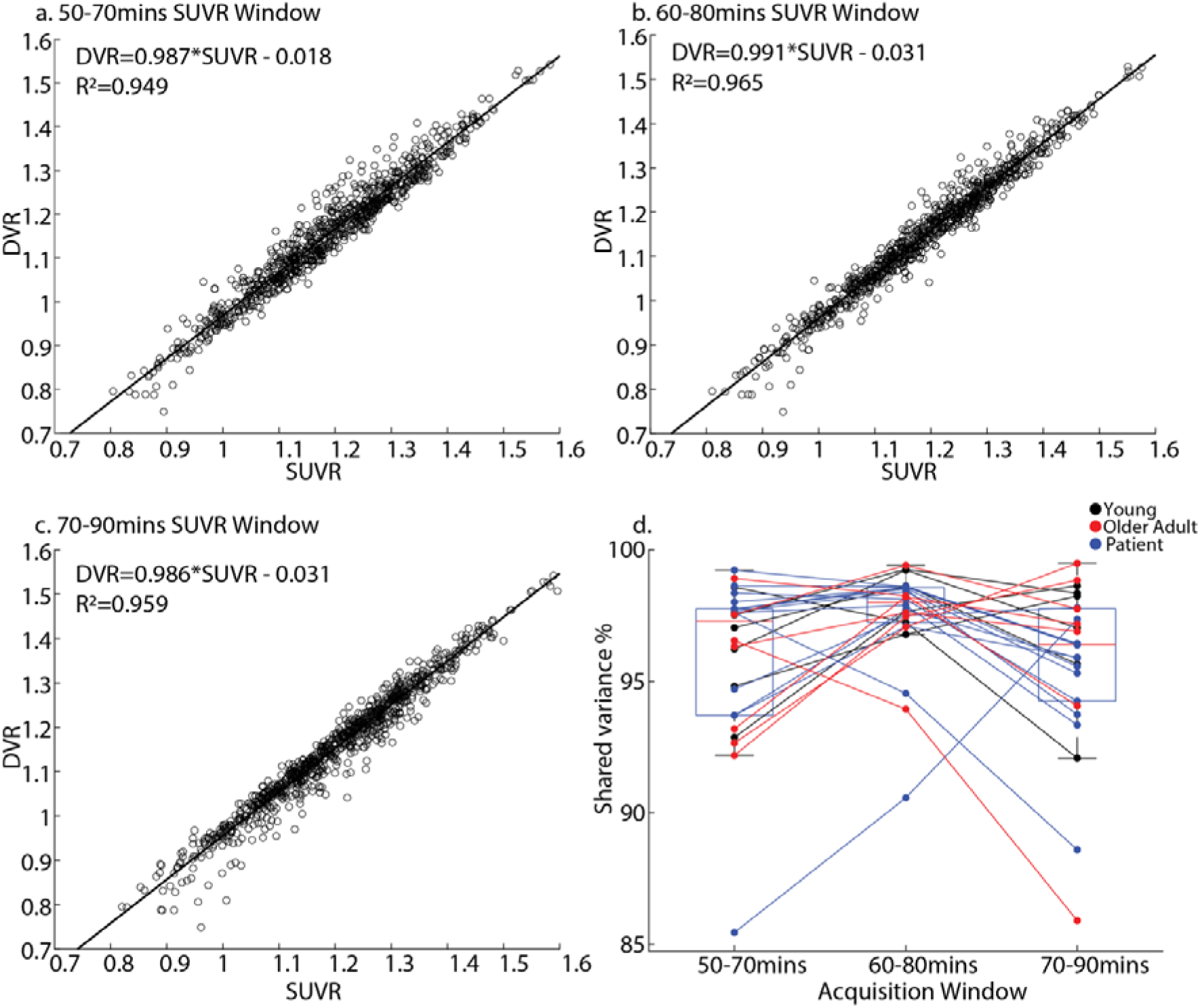
Optimal window for SUVR derivation. Scatter plots show the relationship between SUVR and DVR in each cortical ROI for each subject using an acquisition window (a.) 50-70 minutes, (b.) 60-80 minutes, or (c.) 70-90 minutes post injection. Text in each panel shows the equation of the least squares fit of SUVR to DVR as well as the shared variance (R^2^) between the measures. The box chart (d.) shows the percent within subject shared variance between SUVR and DVR within cortical ROIs for the different windows used for SUVR derivation.

### Differentiating diagnostic groups using [^18^F]-SynVesT-1 DVR and SUVR

We observed differences in the average uptake across cortical regions between the three groups for both DVR and SUVR data, with the highest uptake in Y, intermediate uptake in O, and the lowest uptake in P (one-way ANOVA: meta-temporal ROI DVR: F(2,25)=6.8, p=0.004, SUVR: F(2,23)=8.57, p=0.0017; temporal lobe DVR: F(2,25)=9.21, p=0.004, SUVR: F(2,23)=10.14, p=0.0007; parietal lobe DVR: F(2,25)=13.91, p<0.001, SUVR: F(2,23)=12.31, p=0.0002; frontal lobe DVR: F(2,25)=11.16, p=0.0003, SUVR: F(2,23)=10.52, p=0.0006) (post-hoc contrasts show in **Figure 4**). Visual inspection of whole brain images also supports the results across specific ROIs (**Supplementary Figure 2**). After applying two tissue compartment PVC to account for potential biases due to differences in grey matter volume, we observed an attenuated difference between Y and O, but P continued to have lower uptake than O (two sample t-test: meta-temporal ROI t(19)=-2, p=0.06; temporal lobe t(19)=-2.2, p=0.04; parietal lobe t(19)=-2.34, p=0.03; frontal lobe t(19)=-1.896, p=0.073) (**Supplementary Figure 3**). Taken together, this suggests there are differences in [^18^F]-SynVesT-1 uptake across the cortex between cognitively normal O and P and that this difference is unlikely to be driven solely by differences in grey matter volume between the groups.

**Figure 4.**
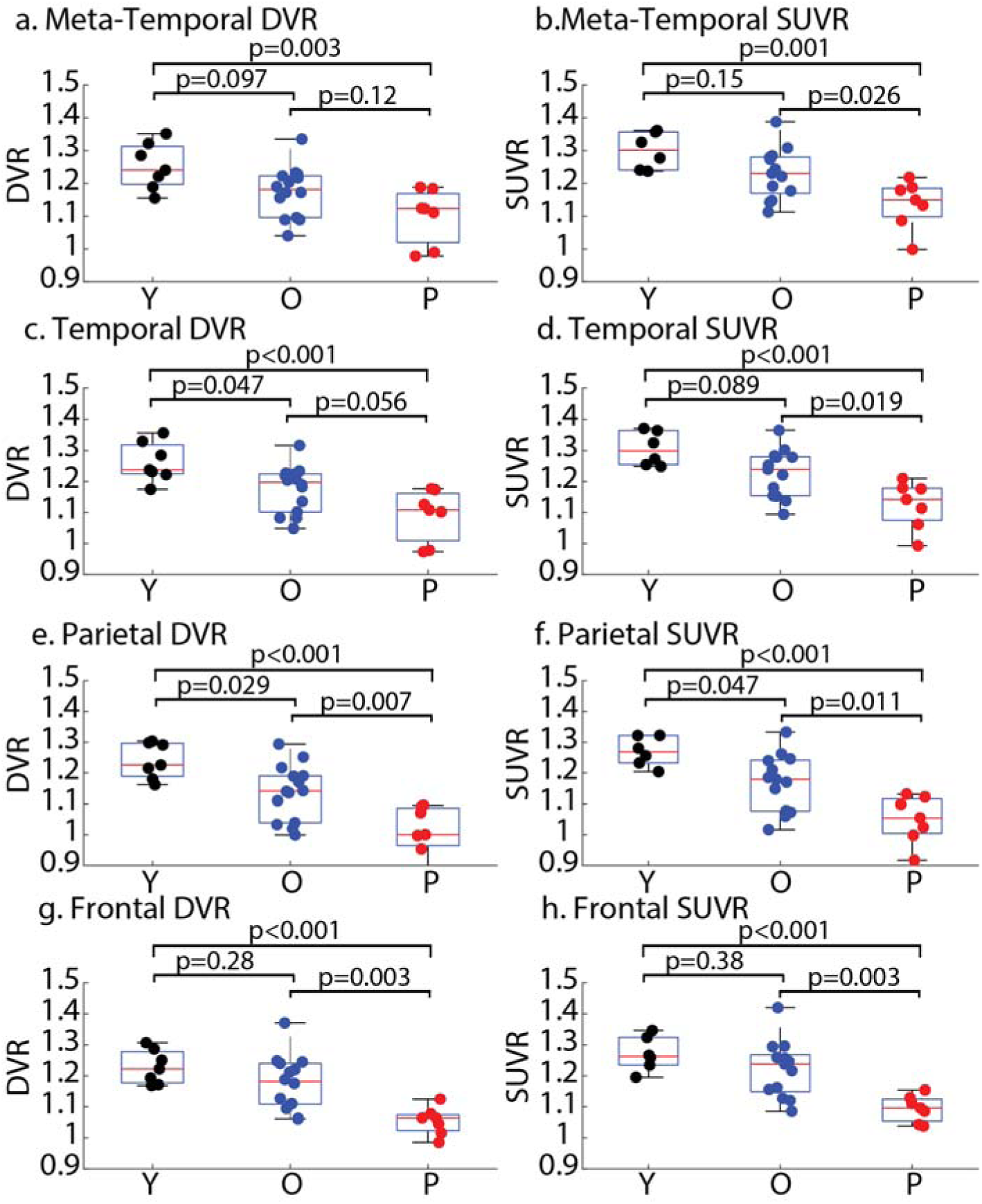
Group differences in DVR and SUVR uptake. Group differences in DVR in the (a.) meta-temporal ROI, (c.) temporal lobes, (e.) parietal lobes, and (g.) frontal lobes. Group differences in SUVR in the (b.) meta-temporal ROI, (d.) temporal lobes, (f.) parietal lobes, and (h.) frontal lobes. Two older adults did not have PET acquisition within the 60-80 minute window and a SUVR value could not be derived. P values for the group wise contrasts within each ROI are shown above the plots.

### [^18^F]-SynVesT-1 uptake mirrors phenotypic tau uptake, tracks A**b** levels, and interacts with grey matter volume when estimating cognition

To assess how tau PET binding related to [^18^F]-SynVesT-1 PET binding we extracted from each participants pair of tau and [^18^F]-SynVesT-1 PET images the uptake across 68 bilateral cortical ROIs. We then correlated (Pearson’s) these two vectors for each subject, whereby a negative correlation coefficient for a participant indicates lower [^18^F]-SynVesT-1 uptake in corresponding regions with high tau PET uptake. Across the combined O and P groups we observed that the correlation values between imaging pairs were significantly less than 0 (t(20)=-3.5,p=0.002) indicating a negative association between the regional binding of tau and [^18^F]-SynVesT-1 PET. Further, we observed that patients had a significantly greater negative association between tau PET and [^18^F]-SynVesT-1 uptake than older adults (t(19)=-4.76, p<0.001), that is, those with a phenotypic spatial distribution of increased tau PET show a similar spatial distribution of decreased [^18^F]-SynVesT-1 PET uptake. Quantitatively this indicates a mirroring of [^18^F]-SynVesT-1 uptake and tau PET uptake in clinical AD phenotypes (**Figure 5**). Qualitative visual inspection of the spatial distributions of [^18^F]-SynVesT-1 uptake and tau PET confirms this mirroring of signal, whereby areas of high tau PET uptake show low [^18^F]-SynVesT-1 uptake. In patients with amnestic dementia due to AD, areas of high bilateral temporoparietal tau PET tracer uptake had lower levels of [^18^F]-SynVesT-1 uptake. The patient with lvPPA due to AD had high tau PET uptake in left temporal cortex, posterior perisylvian, and parietal regions that corresponded to low uptake of [^18^F]-SynVesT-1. Finally, we observed a similar mirroring in the patient with PCA due to AD who had high right > left parieto-occipital tau binding that corresponded with low [^18^F]- SynVesT-1 uptake (**Figure 5**).

**Figure 5.**
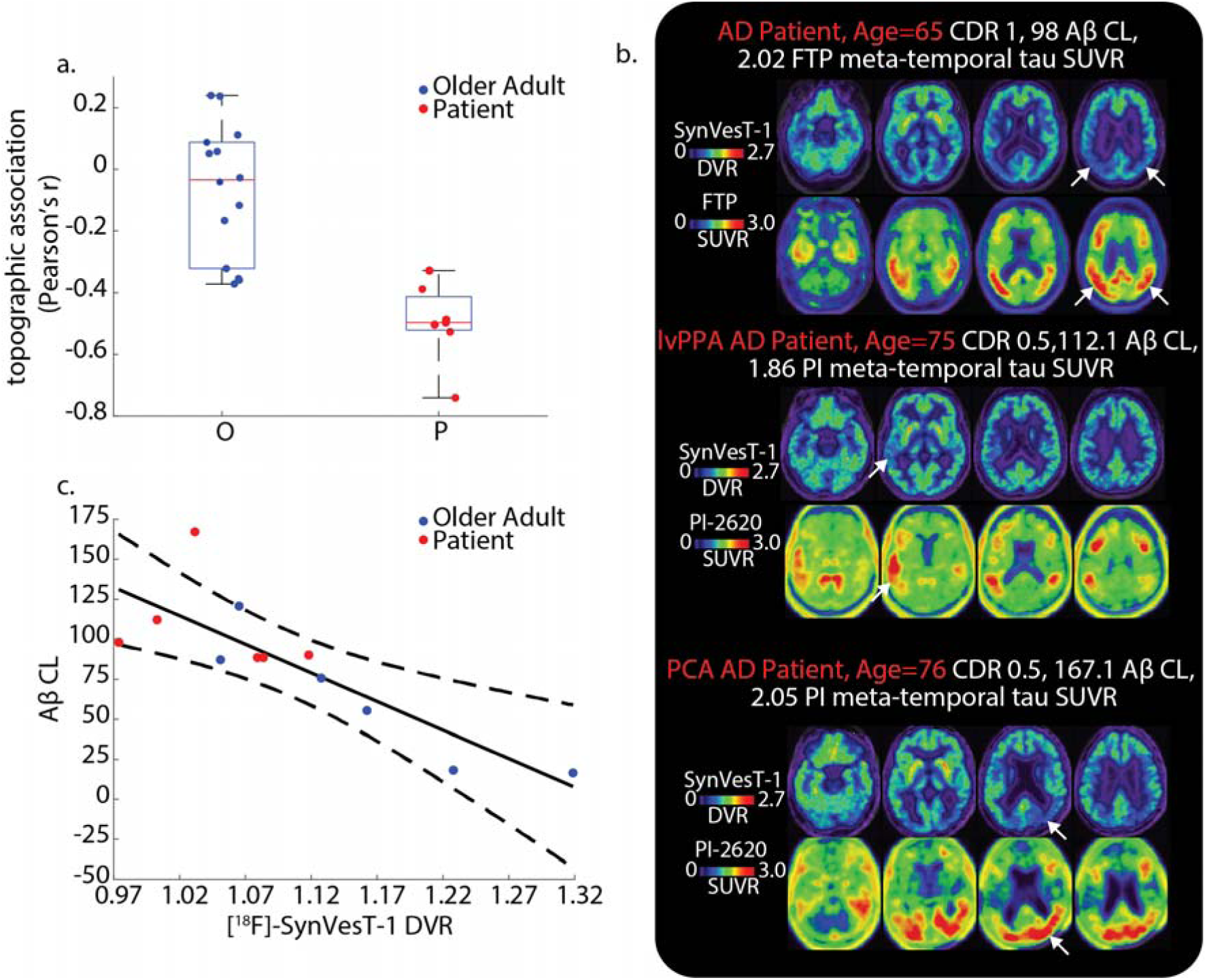
Relationship between [^18^F]-SynVesT-1 uptake and AD pathology. (a) topographic association (Pearson’s correlation coefficient) between [^18^F]-SynVesT-1 uptake and tau-PET uptake for each participant. Negative values indicate regions with higher tau PET uptake show lower [^18^F]-SynVesT-1 uptake for that participant. (b.) [^18^F]-SynVesT-1 and tau-PET binding patterns for three patients with typical amnestic AD (top), lvPPA due to AD (middle), or PCA due to AD (bottom). White arrows show regions of high tau PET uptake (lower panel) and corresponding regions of low [^18^F]-SynVesT-1 uptake (upper panel). (.c) Association between Ab CL and cortical [^18^F]-SynVesT-1 uptake. Black solid line is the least squares fit between Ab CL and [^18^F]-SynVesT-1, dashed black lines are the 95% confidence intervals of this fit.

Next, we tested whether the cortical uptake of [^18^F]-SynVesT-1 is correlated with Ab CL level in the 12 Ab participants with a visually positive scan (6 P, 6 O). We observed a strong negative correlation between CL levels of Ab and average cortical [^18^F]-SynVesT-1 uptake without and with PVC (Pearson’s correlation: without PVC r(10)=-0.83, p<0.001; with PVC r(10)=-0.68, p=0.016), suggesting that the level of Ab may impact global SV2A density (**Figure 5**). A multiple regression including the interaction between clinical diagnosis and [^18^F]-SynVesT-1 uptake showed no significant interaction or main effects with diagnosis on Ab CL when including [^18^F]-SynVesT-1 uptake (Adj R^2^=0.61, p=0.014; [^18^F]-SynVesT-1 t=- 3.19 p=0.013; Diagnosis t=-0.66 p=0.53; [^18^F]-SynVesT-1*Diagnosis t=0.69. p=0.51) (**Supplementary Figure 4**). Significance of regression coefficients was the same using PVC [^18^F]-SynVesT-1 uptake.

Using a multiple regression for 27 participants, including age as a confounding variable, we modelled effects of meta-temporal ROI [^18^F]-SynVesT-1 uptake, meta-temporal ROI grey matter volume (normalised by total intracranial volume), and their interaction on MMSE. One patient with a very low MMSE score was omitted from this analysis (MMSE=4). This model explained substantial variance in MMSE (adjusted R^2^=0.6, p<0.001), with significant main and interaction effects (meta-temporal ROI volume t=2.60, p=0.016; meta-temporal ROI [^18^F]-SynVesT-1 uptake t=2.55, p=0.018; meta-temporal ROI [^18^F]-SynVesT-1 uptake*meta-temporal ROI volume t=-2.27, p=0.034) (**Figure 6**). Individuals with low grey matter volume but high [^18^F]-SynVesT-1 uptake had relatively preserved cognition (**Figure 6a**). This result was the same when modelling the interaction using PVC meta-temporal ROI [^18^F]-SynVesT-1 uptake (adjusted R^2^=0.58, p<0.001) with significant main and interaction effects (meta-temporal ROI volume t=2.41, p=0.025; meta-temporal ROI [^18^F]-SynVesT-1 uptake t=2.33, p=0.03; meta-temporal ROI [^18^F]-SynVesT-1 uptake*meta-temporal ROI volume t=-2.06, p=0.05) (**Figure 6b**). Model interpretation and results were the same when excluding the Y participants, although statistical power is greatly reduced (**Supplementary Figure 5**). This further supports the additive information of [^18^F]-SynVesT-1 PET relative to MRI atrophy markers and suggests that increased density of SV2A may offset the effects of lower cortical volume on cognition.

**Figure 6.**
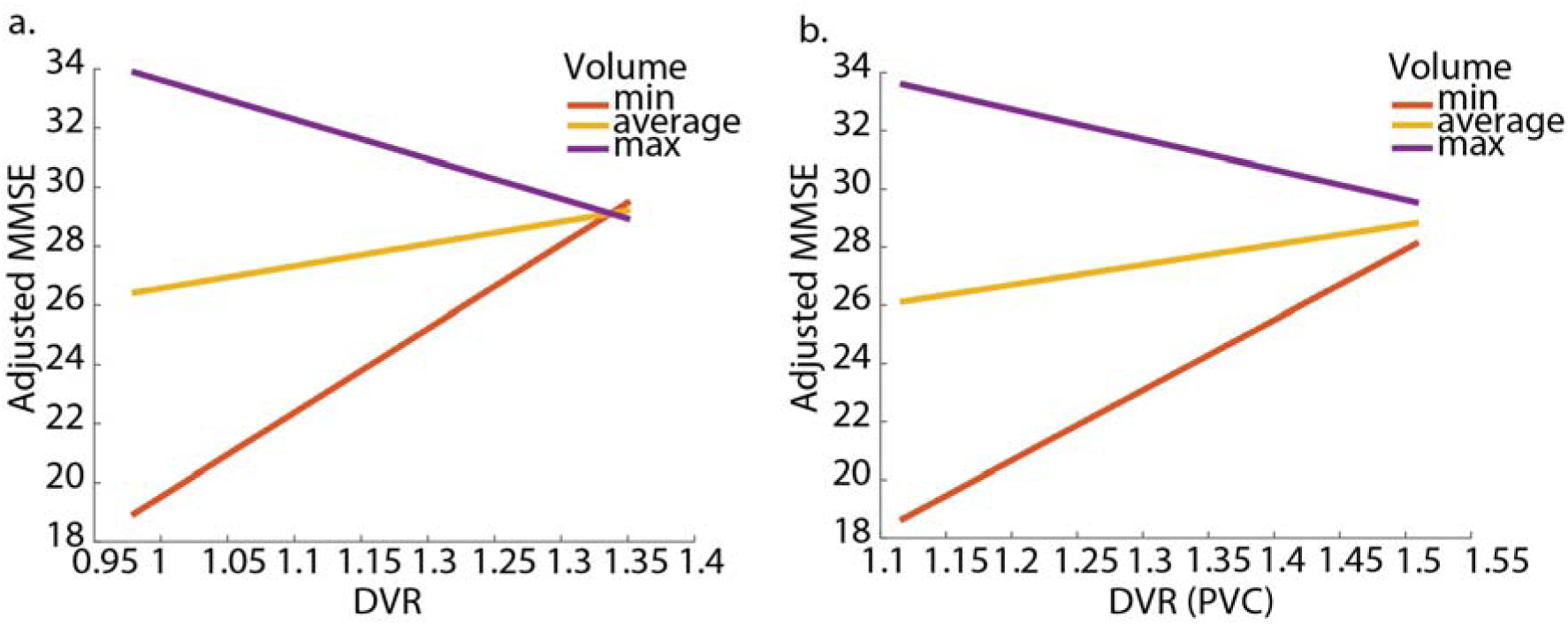
Interaction between average meta-temporal ROIs [^18^F]-SynVesT-1 and volume predicts cognition. Interaction between average meta-temporal ROI volume and [^18^F]-SynVesT-1 uptake in predicting MMSE using (a.) non-PVC and (b.) PVC SynVesT-1 DVR. Individual lines indicate predicted MMSE at different level of meta-temporal ROI [^18^F]-SynVesT-1 uptake for three different levels of meta-temporal ROI volume: the minimum volume in the sample (red), the average volume in the sample (yellow) and the maximum volume in the sample (purple). MMSE values on y-axes are adjusted for age and marginal levels are not restricted to the ceiling of the MMSE, as such the marginal level shown at the maximum cortical volume exceeds the ceiling of un-adjusted MMSE scores (30).

## Discussion

Here we investigated variation in the uptake of [^18^F]-SynVesT-1 PET in a heterogenous sample comprised of young and cognitively older adults as well as patients with varied AD phenotypic presentations. Within this sample we undertook a range of PET modelling optimisation procedures to determine the optimal reference tissue to characterise *in vivo* synaptic density in a diverse dataset. Finally, we investigated the variation in [^18^F]-SynVesT- 1 PET uptake across ageing and AD phenotypes and how this relates to AD biomarkers and cognitive impairment.

Using reference tissue modelling approaches we show that the cerebellar grey matter provided the optimal characteristics for a reference region, fitting observed TACs best across age and diagnostic categories. To date, most SV2A PET studies using reference tissue modelling have used the centrum semiovale as a reference region (*39*). Recently however a cerebellar reference region has been used, particularly when investigating synaptic changes in AD (*28*,*31*,*36*,*52*). Here, we show that for reference tissue modelling the cerebellar grey matter works better than an eroded white matter reference region encompassing the centrum semiovale. Initial reference region selection for SV2A PET using [^11^C]-UCB-J determined the centrum semiovale as a preferred reference region (*2*). Subsequent tracer blocking studies using the antiepileptic drug padsevonil supported this region, showing displaceable binding in cortical, subcortical, and cerebellar regions, but no significant difference in centrum semiovale uptake (*37*). However, additional blocking studies using a different anti-epileptic targeting SV2A, levetiracetam, reported a 12-22% reduction in [^11^C]-UCB-J uptake within centrum semiovale highlighting that although there is low uptake within this region, it is not devoid of displaceable binding (*2*,*47*). A subsequent study using [^18^F]-SynVesT-1 PET reproduced this finding, showing that there was displaceable binding within the centrum semiovale (*5*). These blocking studies suggest that neither the cerebellum nor centrum semiovale are perfect reference regions for SV2A PET; however, the cerebellar grey matter may be the preferred reference region based on the consistency of binding across diagnostic groups and the reliability of predictions of TACs using SRTM2.

Although the cerebellar grey matter performs well at reproducing observed TACs, care needs to be taken in the interpretation of values derived from dynamic analysis (i.e. DVR). SRTM2 assumes no displaceable binding in the reference region (*48*). However, as the cerebellar grey matter shows displaceable binding, then terms are no longer cancelled out in the derivation of binding potential (BP) using SRTM2, and as such, the DVR no longer represents BP+1 (*53*). Comparison of the cerebellum and centrum semiovale using gold standard arterial sampling however shows a similar distribution volume across the cortex (*31*). Assumptions regarding non-displaceable binding are not required in the interpretation of SUVR values and given the tight association between DVR and SUVR in the 60-80 minute post-injection acquisition window, this may serve a more appropriate summary image to use in analysis.

We observed that [^18^F]-SynVesT-1 PET uptake is reduced across the cortex in patients with AD with a negative association between the level of Aβ burden and cortical [^18^F]-SynVesT-1 PET uptake when examining cognitively normal and impaired participants together. Furthermore, we show that the spatial extent of tau pathology that is characteristic of AD phenotypes is mirrored by reduced [^18^F]-SynVesT-1 PET uptake. This extends previous work in typical AD showing that SV2A PET decreases are associated with both Aβ (*28*,*29*) and tau PET increases (*28*,*30*,*34*,*36*) and that longitudinal reductions in SV2A PET follows a similar Braak staging scheme as tau PET (*32*). Further, after accounting for grey matter differences in [^18^F]-SynVesT-1 PET uptake using PVC, we did not observe significant differences in cortical synaptic density between young and cognitively normal older adults, but significant differences between the normal adults and patients persisted. The lack of association between SV2A PET in cortical regions and aging replicates previous imaging studies (*8–10*).

Next, we observed an interaction between meta-temporal [^18^F]-SynVesT-1 PET uptake and volume when predicting cognition whereby individuals with low meta-temporal volume but elevated [^18^F]-SynVesT-1 PET uptake show preserved MMSE scores. This suggests that there is additive information between MRI measures of neurodegeneration and [^18^F]- SynVesT-1 PET. Previous work has shown significant differences in SV2A PET between AD patients and controls (*26*,*28*,*31*,*32*,*34–36*,*52*) as well as associations between temporal SV2A PET uptake and cognition (*26*,*52*). When incorporating both volume and SV2A PET in the same models, prior work either controlled for grey matter differences through PVC or compared relative effect sizes of SV2A PET and grey matter volume when predicting cognition. Here we extend these findings highlighting the interactive effects between grey matter volume and [^18^F]-SynVesT-1 PET uptake on preserved cognition. This finding may be suggestive that maintaining higher synaptic density throughout neurodegenerative processes is a possible mechanism and *in vivo* marker of resilience.

There are several limitations to this work. First, the sample size was small within each group, and patients were younger than older controls. However, our findings showing lower [^18^F]- SynVesT-1 PET uptake in patients was significant compared to both young and cognitively normal older adults giving us confidence that the disease related reduction in presynaptic density we observed is greater than that seen from age alone. Furthermore, our sample of patients had substantial heterogeneity but given the sample size limitations we could only investigate broad differences between older adults and patients. Second, we used a simple two tissue compartment PVC approach. Several PVC approaches have been applied to SV2A PET (*38*), with comparisons between alternative PVC indicating an iterative yang approach may be best for [^11^C]-UCB-J (*54*). However, our findings show no significant age or group variation in uptake within the eroded white matter, which supports collapsing the white and grey matter into a single component. Finally, the underlying meaning of variable SV2A PET uptake is not perfectly understood. An increased number of SV2A binding sites may represent a true increase in the number of synapses in a given region thus representing increased synaptic density. Alternatively, increased SV2A binding sites may be due to an increased number of transmembrane sites on presynaptic vesicles, or a higher number of presynaptic vesicles, neither of which necessitate increased synaptic density (*55*). Prior autoradiography findings do find an association between SV2A and an alternative marker of synaptic density, synaptophysin (*2*), but further work is required to unequivocally determine that variation in SV2A PET binding captures variation in synaptic density.

## Conclusions

Tracking the alterations of synapses throughout neurodegenerative processes offers new insights into the concomitant pathophysiology of AD as well as putative mechanisms that may preserve cognition in the face of neurodegeneration. Here, we present a modelling study of [^18^F]-SynVesT-1 PET paired with an investigation of SV2A variations across ageing and AD clinical phenotypes. Using a reference region taken from the cerebellar grey matter and a truncated PET acquisition window of 60-80 mins post injection to derive SUVR images we make modelling suggestions that may facilitate a broader use of [^18^F]-SynVesT-1 PET in clinical research. We next show using dynamic [^18^F]-SynVesT-1 PET there is a close negative association between the pattern of [^18^F]-SynVesT-1 and tau PET in patients with varied phenotypic presentations of AD. Finally, we observed interactive relationships between preserved SV2A density and neurodegeneration whereby individuals with lower meta-temporal grey matter volume but higher meta-temporal [^18^F]-SynVesT-1 PET binding showed preserved cognition, a possible indicator of resilience.

## Key Points

QUESTION: Determine the optimal approach to derive reference tissue images of [^18^F]- SynVesT-1 PET and assess if these images capture differences in synaptic density in varied Alzheimer’s Disease phenotypes.

PERTINENT FINDINGS: In this cohort study we found that a reference region from the cerebellar grey matter was significantly better at modelling ground truth [^18^F]-SynVesT-1 PET time activity curves using reference tissue modelling. Using whole brain [^18^F]-SynVesT-1 PET ratio images we found that regions with high tau PET uptake had lower synaptic density. Finally, we found a significant interaction between grey matter volume and [^18^F]- SynVesT-1 PET uptake, whereby individuals with low grey matter volume but high synaptic density showed better cognitive performance.

IMPLICATIONS FOR PATIENT CARE: These findings provide guidelines into the optimal modelling of [^18^F]-SynVesT-1 PET and highlights the clinical significance of [^18^F]-SynVesT-1 PET in understanding the pathology of Alzheimer’s Disease.

