## supplementary information for "Imaging synaptic density in ageing and Alzheimer’s Disease with [^18^F]-SynVesT-1"

**Supplementary Tables and Figures:**

| **Measure** | **Cerebellar Grey** | **Whole Cerebellum** | **Eroded white matter** |
| --- | --- | --- | --- |
| Mean SSE | 356716 | 423020 | 496344 |
| SSE CV | 0.41 | 0.41 | 0.47 |
| Mean k_2_’ | 0.019 | 0.021 | 0.027 |
| k_2_’ CV | 0.08 | 0.08 | 0.09 |

**Supplementary Table 1. SRTM2 results**. Mean Sum Squared Error (SSE) is calculated from the average error of the SRTM2 derived time activity curves for the 34 bilateral cortical Desikan-Killiany ROIs. The k2’ used throughout is the average value of that produced the lowest error in fitting the cortical data for each subject. Variability for each parameter/metric is shown as the coefficient of variation (CV).

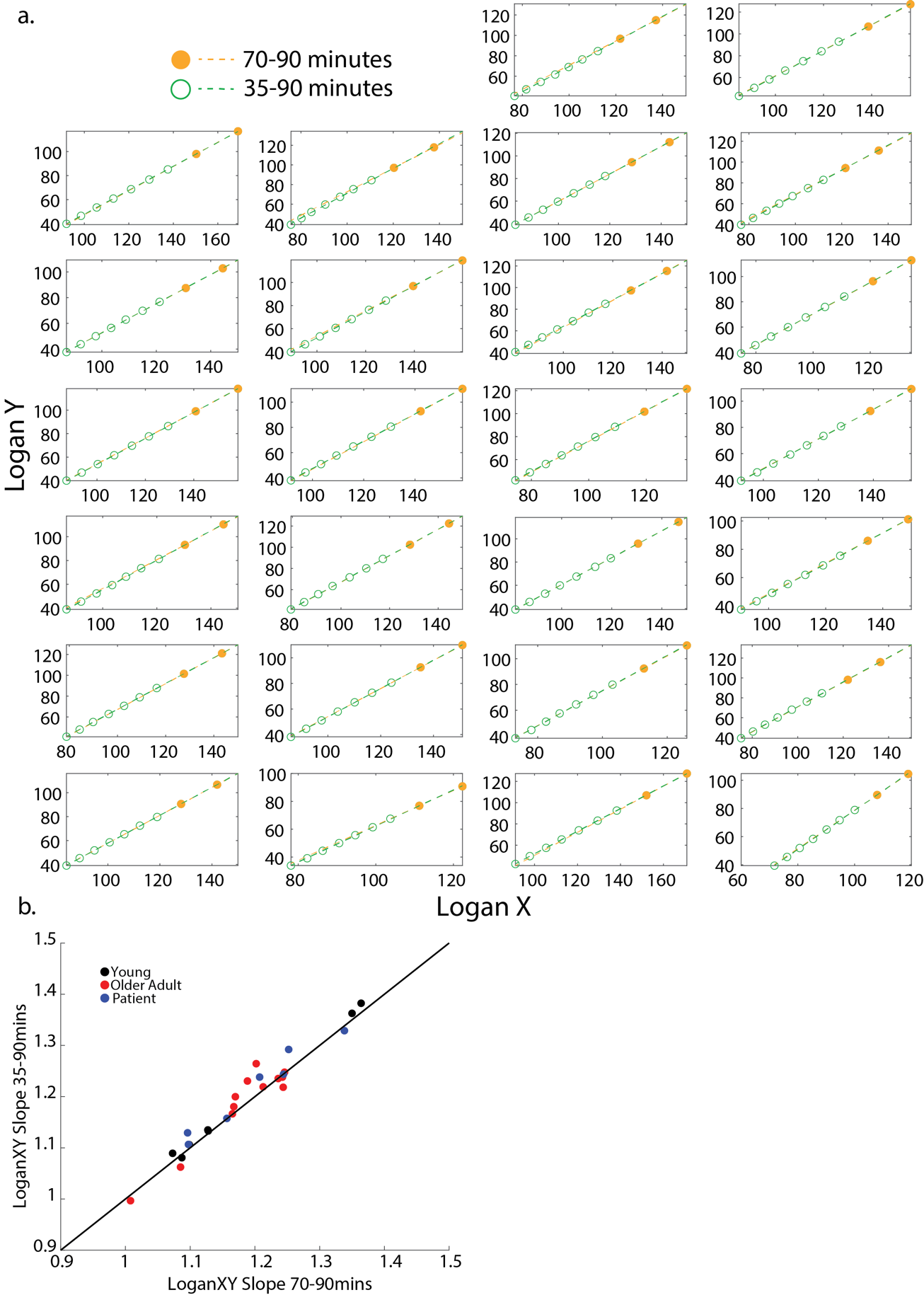

**Supplementary Figure 1. Stability of slope of Logan graphical analysis.** (a.) Logan X vs. Logan Y for each participant. Green circles are the values at each PET frame from 35 - 90 minutes, orange circles are the values for frames 70-90 minutes. Dashed green line is the slope of Logan X vs. Logan Y across the full 35-90 minute window, dashed orange line is the slope of Logan X vs. Logan Y for the 70-90minute window. (b.) relationship between the 35-90 minute slope (y axis) and the 70-90 minute slope. Black line is the identity line (i.e. y=x).

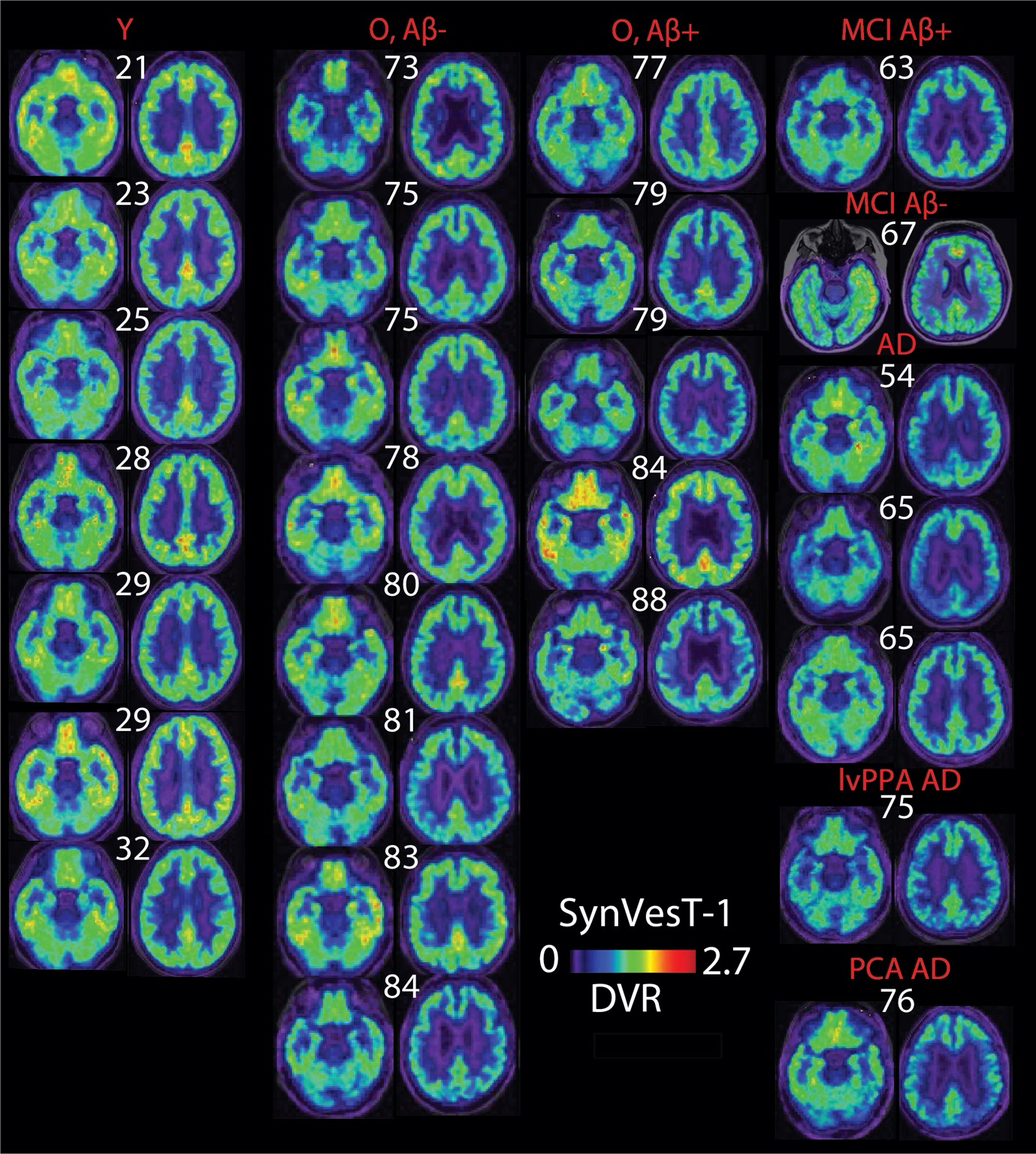

**Supplementary Figure 2. Whole brain [^18^F]-SynVesT-1 DVR images.** Representative slices of the whole brain DVR images for young adults (Y), older adults (O), and patients (P). Red titles represent the group or AD phenotype for the images below. White numbers above brain slices indicate the participant’s age. All images are scaled to the same minimum (0) and maximum (2.7) DVR values. Alzheimer’s Disease (AD), Mild Cognitive Impairment (MCI), Aβ negative (-), Aβ positive (+), posterior cortical atrophy (PCA), logopenic variant primary progressive aphasia (lvPPA). MCI Aβ+ patient carries the *presenilin 1* (*PSEN1*) mutation.

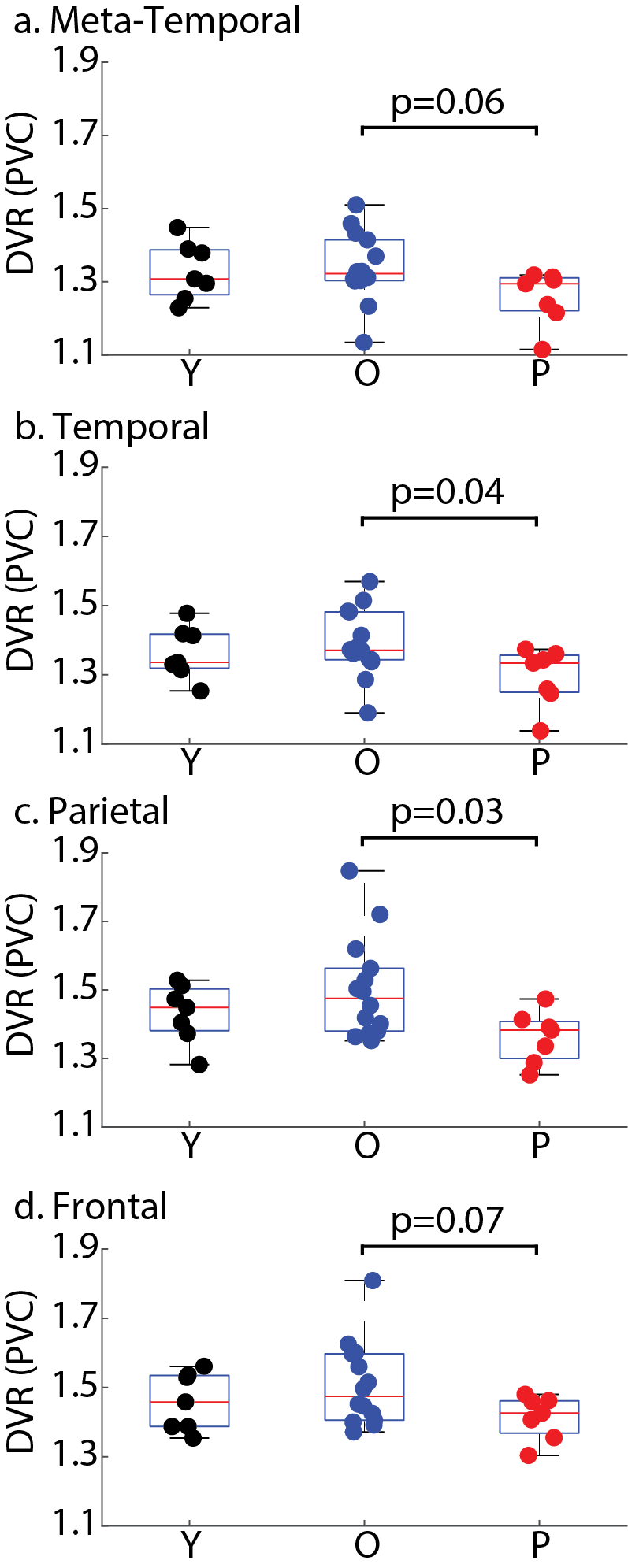

**Supplementary Figure 3. Group differences in PVC [^18^F]-SynVesT-1 DVR.** Group differences in PVC [^18^F]-SynVesT-1 DVR in the (a.) meta-temporal ROI, (b.) temporal lobes, (c.) parietal lobes, and (d.) frontal lobes. P values for the group wise contrasts of older adults (O) and patients (P) within each ROI are shown above the plots.

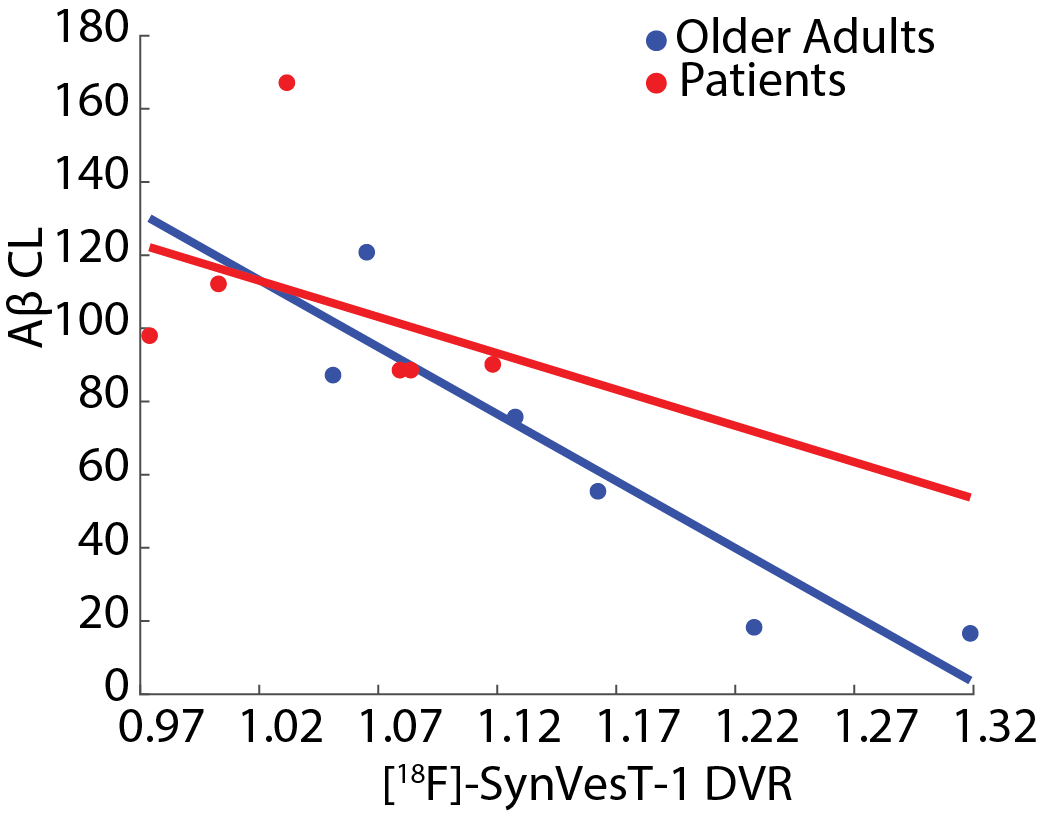

**Supplementary Figure 4. Association between Aβ Centiloid (CL) and cortical [^18^F]-SynVesT-1 uptake (DVR).** Red and blue lines show the least squares fit between Aβ CL and [^18^F]-SynVesT-1 DVR for patients and older adults respectively.

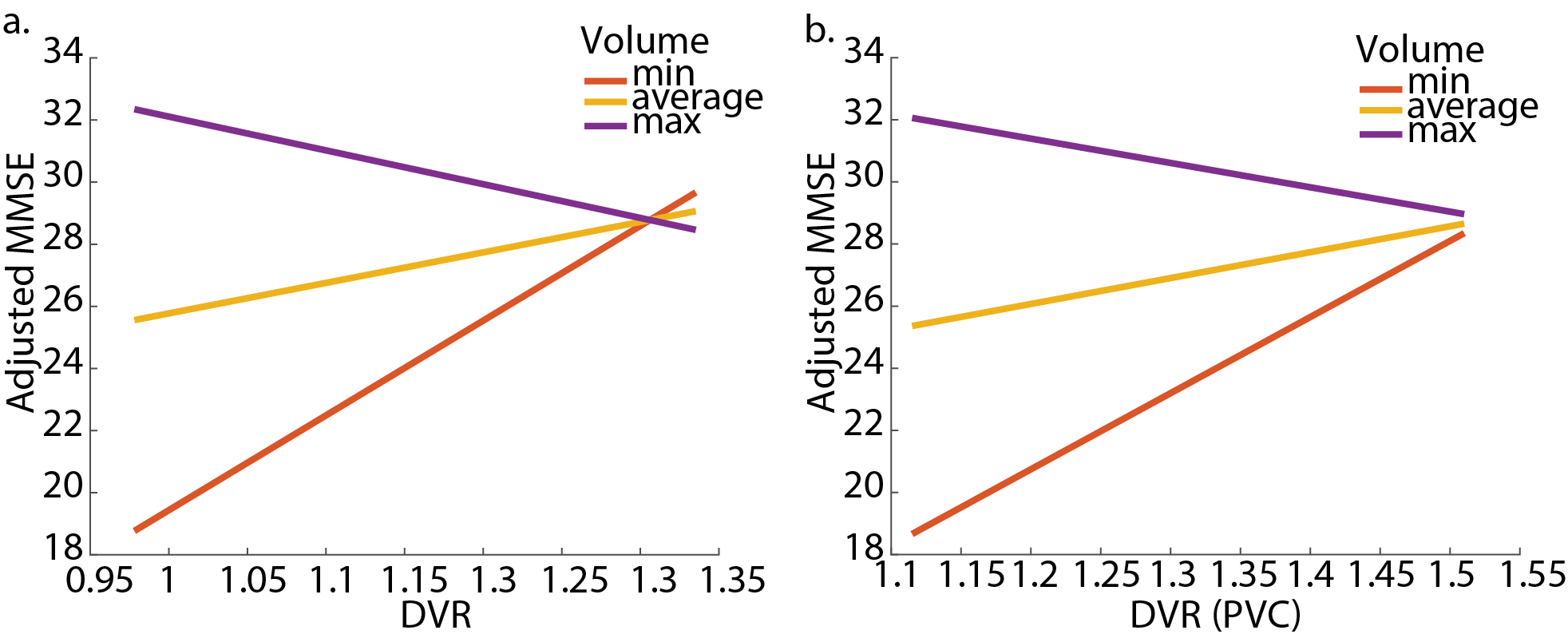

**Supplementary Figure 5. Interaction between average meta-temporal ROI [^18^F]-SynVesT-1 uptake and volume predicts cognition in older healthy adults and patients.** Interaction between average meta-temporal ROI [^18^F]-SynVesT-1 uptake and volume in predicting MMSE using (a.) non-PVC and (b.) PVC [^18^F]-SynVesT-1 DVR. Individual lines indicate predicted MMSE at different level of meta-temporal ROI [^18^F]-SynVesT-1 uptake for three different levels of meta-temporal ROI volume: the minimum volume in the sample (red), the average volume in the sample (yellow) and the maximum volume in the sample (purple).
